# On species delimitation, hybridization and population structure of cassava whitefly in Africa

**DOI:** 10.1101/836072

**Authors:** S Elfekih, WT Tay, A Polaszek, KHJ Gordon, D Kunz, S Macfadyen, TK Walsh, S Vyskočilová, J Colvin, PJ De Barro

## Abstract

The *Bemisia* cassava whitefly complex includes species that cause severe crop damage through vectoring cassava viruses in eastern Africa. Currently, the cassava whitefly complex is divided into species and subgroups based on very limited molecular markers that did not allow clear definition of species and population structure. Based on 14,358 genome-wide SNPs from 63 cassava whitefly individuals belonging to sub-Saharan African (SSA1, SSA2 and SSA4) species, and using a well-curated mtCOI gene database, we show clear incongruities in previous taxonomic approaches underpinned by effects from pseudogenes. We show that the SSA4 species is part of the SSA2 species, and that populations of the SSA1 species comprise of south-eastern (Madagascar, Tanzania) and north-western (Nigeria, Democratic Republic of Congo and Burundi) sub-species that show signatures of allopatric incipient speciation, with a hybrid zone separating these adjacent sub-species. These findings provide the first genomic insights into the evolution and molecular ecology of a highly cryptic hemipteran insect complex in African, and allow the systematic use of genomic data in management and control strategies for this important cassava pest.

## Introduction

Confidence in species identification underpins strategies for biological conservation and management of natural resources, providing insight into the evolutionary forces that drive species diversification. While morphological differences have enabled many species to be identified, cryptic species complexes present a particular challenge due to their general lack of readily distinguishable morphological characters. Collecting ecological and life history data, including host plant use and mating behaviour, can be used effectively to mitigate and address these challenges (Liu, et al. 2007; Vyskočilová, et al. 2019; Vyskočilová, et al. 2018), but are often time-consuming and impractical, especially with large numbers of samples. Incorporating DNA markers, i.e., diagnostic mitochondrial DNA (mtDNA) markers (e.g., Behere, et al. 2008; Elfekih, et al. 2018b; Walsh, et al. 2019), genome-wide sequencing (Anderson, et al. 2016; Elfekih, et al. 2018a), and whole-genome resequencing (Anderson, et al. 2018), can increase confidence in differentiating between cryptic species in a complex and identifying inter-species hybrids.

The challenges can become critical when individual cryptic species in a complex have a serious impact on health or agricultural productivity. An example of the latter is presented by insect vectors of virus diseases affecting cassava, the most important food crop grown in Africa (55.2% global production share; average from 1994-2011; FAOSTAT 2017). Since the 1990’s, cassava production has been affected by a pandemic of cassava mosaic disease (CMD) caused by a number of cassava mosaic geminiviruses (CMGs) (Legg, et al. 2014; Patil and Fauquet 2009). Although cassava is also widely grown in Latin America (its original home) and South East Asia, CMD has been reported in Africa since the early 1970’s (Macfadyen, et al. 2018) and more recently in South-East Asia (Minato, et al. 2019; Wang, et al. 2016). While the vector responsible for the spread of the CMD in South-East Asia is not fully understood, the epidemic in east and central Africa was associated with the distribution of invasive Sub-Saharan African (SSA) species of the *Bemisia* whitefly complex (Legg, et al. 2014). Invasive whitefly species have become major pests of global agriculture, spreading plant virus diseases and becoming resistant to chemical control agents (De Barro, et al. 2011). In particular, the invasive cassava whitefly lineage SSA2 (previously ‘Ug2’; Boykin, et al. 2017) has been found in regions affected by the CMD pandemic, and since 2007 has also successfully expanded into southern Europe (Hadjistylli, et al. 2015), while another lineage, the SSA1 (previously ‘Ug1’; Boykin, et al. 2017), was found beyond the pandemic-affected geographical regions (i.e., widespread throughout the African continent).

Defining the actual CMD vectors in Africa remains a challenge due to the cryptic nature of the many species in the Aleyrodidae whitefly *Bemisia* genus (De Barro, et al. 2011; Lee, et al. 2013; Mugerwa, et al. 2018; Vyskočilová, et al. 2018). At least 38 cryptic species have now been recognised within what was previously considered a single *Bemisia tabaci* (Gennadius) species (Martin and Mound 2007) through the characterisation of a partial mitochondrial DNA cytochrome oxidase subunit I (mtCOI) 3’ gene region of 657bp (De Barro, et al. 2011; Dinsdale, et al. 2010; Kunz, et al. 2019b). However, re-analyses of partial mtCOI sequences in GenBank relating to the Mediterranean (MED) species by Vyskočilová et al. (2018), and of the *B. tabaci* standard partial mtCOI dataset (Boykin, et al. 2017) by Kunz et al. (2019b), demonstrated that a substantial portion (*ca*. 60-65%) were artefacts of nuclear mitochondrial DNA sequences (NUMTs) or other PCR artefacts (Elfekih, et al. 2018a; Kunz, et al. 2019b; Tay, et al. 2017a). This analysis, together with molecular characterisation of the original voucher specimens, is beginning to disentangle the species complex, identifying the Mediterranean (MED) species as the true *B. tabaci* (Tay, et al. 2012), the *B. tabaci* AsiaII_7 species as *B. emiliae* Corbett (Tay, et al. 2017b), assisting with identifying historical species ranges (Kunz, et al. 2019a; Tay, et al. 2017b; Tay, et al. 2012), and provided phylogenetic evidence that the cassava whitefly (i.e., SSA1, SSA2, SSA3, SSA6, SSA9) were ancestral to the ‘*B. tabaci* cryptic species complex’ (Kunz, et al. 2019b). The advent of high-throughput sequencing (HTS) technologies has driven the availability of genome-wide data that is further transforming the *Bemisia* taxonomy. HTS has confirmed that the *B. tabaci* Middle East Asia Minor 2 (MEAM2), previously recognised as a distinct species, is in fact the result of an artefact of sequencing nuclear mitochondrial DNA sequences (NUMTs) (Tay, et al. 2017a; Elfekih, et al. 2018a).

The taxonomy, population structure and evolutionary genetics of the cassava whitefly *Bemisia* species complex (i.e., SSA1, SSA2, SSA3, SSA4, SSA6, and SSA9; see Kunz, et al. 2019b) remain poorly understood, despite the importance of defining and managing the vectors of CMD. In a recent study of taxonomic structure and gene flow among genetically diverse SSA1, SSA2, SSA3 and SSA4 species, Wosula et al. (2017) described serious discrepancies between their results obtained either from mtCOI or by nextRAD sequencing, with possible implications for understanding the dynamics of the CMD-transmitting species. The findings of Wosula et al. (2017) have wider implications to the future usage of this partial mtCOI gene as the current standard DNA marker for species identification of *B. tabaci* and related *Bemisia* cryptic species. This is especially so in the sub-Saharan African *Bemisia* species context, where species identified to-date would need to be reanalysed using a genome-wide sequencing approach. The diversity within genome-wide SNP data, and complex gene flow patterns existed between these newly defined SSA genetic groups (Wosula, et al. 2017) would appeared to also have a different operational taxonomic unit to challenge the traditional ‘species’ unit.

In this study, we seek to understand why the African cassava whitefly SSA1 (which is also further refined into sub-groups 1, 2, and 3 in some studies; e.g., Legg, et al. (2014)), SSA2, and SSA4 species, should have such conflicting species status and admixture patterns depending on whether the mtCOI or nuclear DNA markers were used. To reconcile these contrasting findings, we re-analysed the mtCOI data of Wosula et al. (2017) and compared it to a strictly filtered set of mtCOI sequences for species status delimitation. Identification and removal of NUMT-derived mtCOI resolved conflict of results from mtCOI discrepancies as reported by Wosula et al. (2017). Reanalyses of genome-wide SNP data enabled insights into SSA1-species-wide population substructure to identify the geographic contact zone and signature of intra- and inter-species hybridization. Our findings provide powerful insights into the evolutionary forces shaping incipient speciation and vector competence of this agriculturally invasive species complex with the capacity to impact on food security in sub-Saharan Africa.

## Material and Methods

### mtCOI data handling

The mtCOI sequences from the study by Wosula et al. (2017) were downloaded from GenBank (accession numbers MF417578-MF417602), including all their published sequences and published reference sequences to identify genetic groups within the SSA species. These partial mtCOI sequences were imported into Geneious R11.1 and aligned with reference sequences (Suppl. Fig. 1) using the MAFFT alignment program (Katoh and Standley 2013) with default settings. We examined all sequences for insertion/deletions (indels) as signatures of pseudogenes (NUMTs). Sequences that did not contain indels were assessed also for amino acid substitution patterns at evolutionary conserved regions in the partial COI gene region to assist with further identification of potential NUMTs. Identification of amino acid residue conservation sites across the partial mtCOI gene regions was as described in Kunz et al. (2019b).

### nextRAD data processing

The raw data from nextRAD sequencing of Wosula et al. (2017) was accessed from GenBank (SRP103541). The fastq sequences were screened using FastQC for quality control. The retained reads were trimmed by quality to a total length of 151 bp using Trimmomatic (Bolger, et al. 2014).

### Mapping quality

We mapped the reads from each individual sample to the *B. tabaci* MEAM1 genome (Chen, et al. 2016) using Burrows-Wheeler Aligner (BWA) v.0.7.12 (Li and Durbin 2010). The alignments used the BWA-MEM algorithm with default settings. The SAM files were converted to BAM output using Samtools (Li, et al. 2009). The BAM files were subsequently sorted and indexed, and checked for quality and mapping percentages per scaffold.

### SNP genotyping

SNPs were first called using a *de novo* approach in PyRAD (Eaton 2014). PyRAD is a pipeline designed for RADseq datasets that aims to capture variation at the species/clade level. It allows clustering of highly divergent sequences and takes into consideration indel variation. We also performed SNP calling in PyRAD, using reads mapped to the MEAM1 reference genome (Chen, et al. 2016). Given the low-depth nature of the data, we analysed the data using an additional pipeline. We used the BAM files generated in BWA as input for the program ANGSD (Korneliussen, et al. 2014), which relies on a statistical approach that takes into consideration genotype uncertainty (Fumagalli, et al. 2013).

### Genetic clusters and tests for introgression

Genetic clusters within the dataset that consists of 63 SSA samples were investigated by Principal Component Analysis (PCA) using the SNPRelate R package (Zheng, et al. 2012). We used ADMIXTURE v.1.3.0 (Alexander, et al. 2009) on the 63-sample dataset to identify genetic clusters and estimate the genetic ancestry of the SSA samples. We ran the program with K ancestral clusters varying from 1 to 10 with each ‘K’ value repeated 100 times. A cross-validation test was performed to determine the optimal K value. The program ANGSD v.0.911 (Korneliussen, et al. 2014) was used to run an ABBA-BABA test (D-statistics) to investigate introgression patterns between the 63 samples (SSA1 and SSA2 species) and with the MEAM1 genome as an outgroup. The ABBA-BABA test is based on the comparison of the number of tree topologies of the ABBA and BABA patterns. In this study, the SSA species group patterns we investigated were as defined by Wosula et al. (2017) and consisted of [ESA+CA/SSA2+SSA4/ECA+WA/MEAM1], where ‘ESA’, ‘CA’, ‘ECA’, and ‘WA’ represented East South Africa, Central Africa, East Central Africa, and Western Africa, respectively. We used the MEAM1 species as outgroup for comparison and to infer introgression patterns. We tested the significance of these ABBA/BABA patterns using the corresponding Z-scores, obtained in ANGSD using a jack-knife procedure. An absolute Z-score value of ≥ 3 is used as a cut-off value (Reich, et al. 2009).

### Phylogenetic analyses

The topology of the phylogenetic relationships between individuals /populations/subspecies and species within the dataset (63 SSA samples and three *B. afer* individuals (TZ_CHA1, TZ_CHA2, TZ_KIL1) as the outgroup) was examined using a maximum likelihood (ML) approach. The SSA species were first identified based on the mtCOI marker, then, a phylogeny was reconstructed using the nextRAD genome-wide SNPs excluding samples with low genotype quality to minimize biases that could potentially be introduced by missing data. The phylogenetic reconstruction was carried out in RaxML v.7.2.8 (Stamatakis 2006) using the GTR substitution model and GTRGAMMA as the GAMMA model of rate heterogeneity, with 1,000 bootstrap replications. The phylogenetic clustering obtained in RAxML was used to infer a population ML tree in TreeMix (Pickrell and Pritchard 2012), in order to identify genetic mixing, the history of population splits and admixture patterns. We ran the TreeMix simulations using 0 to 5 migrations events with 100 bootstrap replications.

## Results

### Integrity of African cassava *Bemisia* SSA mtCOI sequences

Of the 11 SSA4 sequences (originally labelled as ‘Sub-Saharan III: Western Africa-Cameroon/Cassava’; see Berry et al. (2004), Table 1), only two (Cam Ayos 1 WO2 and Cam Ayos 2 WO3) did not exhibit any premature stop codons at the amino acid level or have indels at the nucleotide level (Suppl. Fig. 1). Both of these sequences were also much shorter than the remaining nine sequences within the Sub-Saharan III: Western Africa-Cameroon/Cassava clade. Comparison of amino acid composition against 21 high-throughput sequencing-generated mtCOI genes representing 12 evolutionary divergent *Bemisia* cryptic species (Kunz, et al. 2019b), enabled Cam Ayos 2 WO3 to be identified as a potential NUMT sequence. Indels, stop codons, non-conservation of amino acid residues in highly conserved mtCOI region therefore ruled out 10 of the 11 SSA4 sequences as potential NUMTs. The remaining single, short but ‘functional’ Cam Ayos 1 WO2 sequence (AF344246) that clustered with the 10 NUMT-related SSA4 sequences suggested that as a whole, these SSA4 sequences were likely of nuclear genomic origin. The clustering of SSA4 sequences to SSA2 indicates it likely to be of SSA2 nuclear origin, similar to that reported for *B. tabaci* MEAM2 NUMT that originated from the nuclear genome of MEAM1 (Elfekih, et al. 2018a; Tay, et al. 2017a).

**Table 1.**
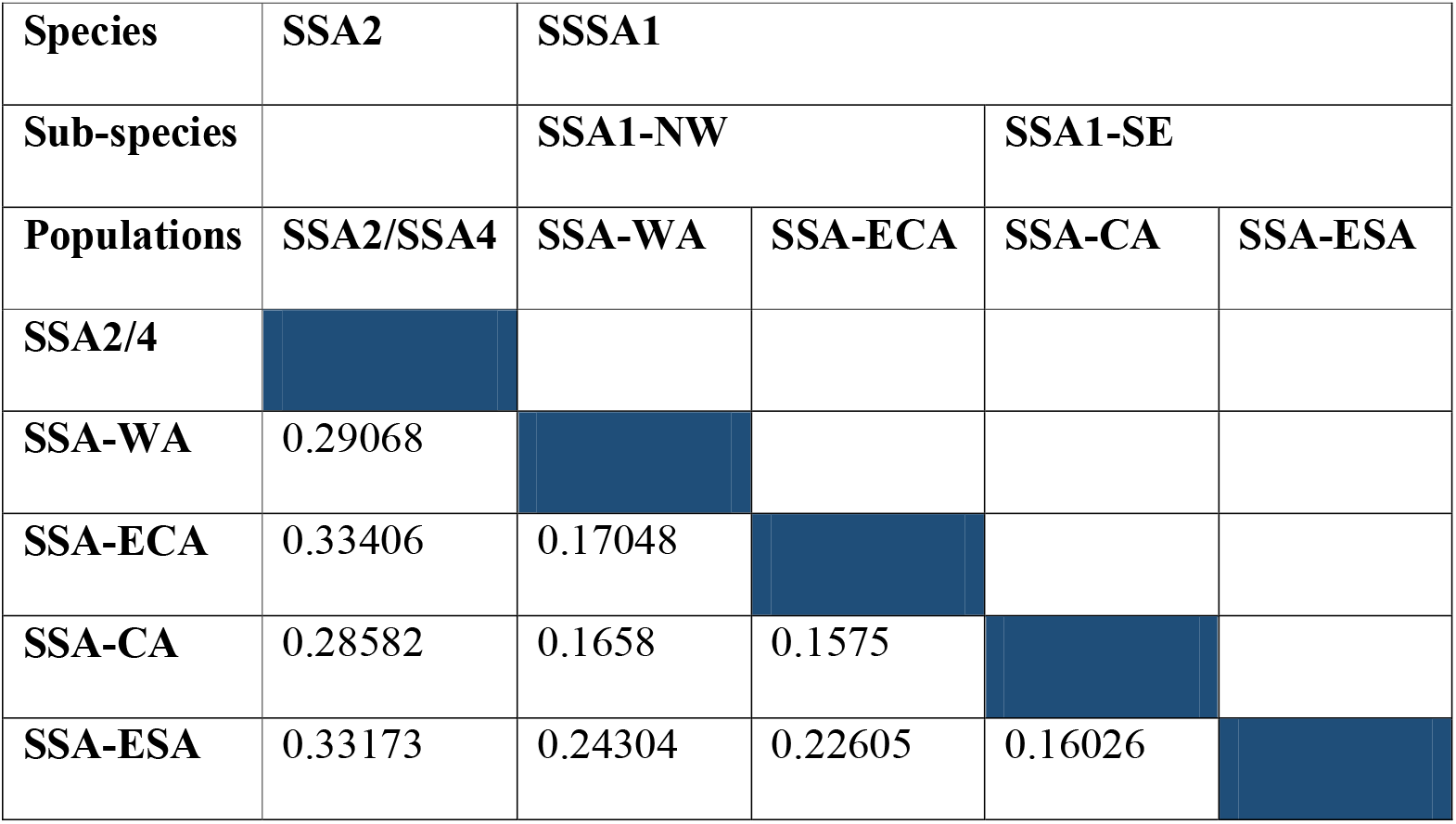
Population substructure based on the *F*-statistics (*F*_st_) of (Weir and Cockerham 1984). Treatment of SSA1 as single species against the SSA2/SSA4 species showed significant population substructure (*F*_st_ range 0.28 – 0.33). This limited gene flow between SSA2/SSA4 and the SSA1 (i.e., SSA-WA, SSA-ECA, SSA-CA. SSA-ESA) supported their separate species status. Treatment of the four SSA1 populations as two well-defined groups, the proposed sub-species, showed that that comprising SSA-WA and SSA-ECA overall shared intermediate levels of gene flow with the SSA-ESA population. Gene flow at the contact zone (SSA-CA) is highest with the SSA-WA, SSA-ECA, and SSA-ESA genetic groups as evident by the low *F*_st_ values.

| Species | SSA2 | SSA1 |  |  |  |
| --- | --- | --- | --- | --- | --- |
| Sub-species |  | SSA1-NW |  | SSA1-SE |  |
| Populations | SSA2/SSA4 | SSA-WA | SSA-ECA | SSA-CA | SSA-ESA |
| SSA2/4 |  |  |  |  |  |
| SSA-WA | 0.29068 |  |  |  |  |
| SSA-ECA | 0.33406 | 0.17048 |  |  |  |
| SSA-CA | 0.28582 | 0.1658 | 0.1575 |  |  |
| SSA-ESA | 0.33173 | 0.24304 | 0.22605 | 0.16026 |  |

The examination of nucleotide substitution patterns identified indels-affected sequences and inconsistencies with respect to number of base substitutions for distinguishing ‘subgroups’ (SG) within the SSA1 species. For example, the SSA1-SG1/SG2 sequences (accession numbers: MF417578-MF417602) were affected by indels that should not be present in a mitochondrial DNA COI protein coding gene, suggesting that these sequences were associated with sequencing errors and/or derived from non-coding NUMT DNA sequences. The re-assessment of the two SSA1-SG5 sequences (MF417585, MF417586) showed that they differed from the SSA1-SG3 sequences by only three C/T substitutions that resulted in 0.56% nucleotide difference. The basis for suggesting MF417585 and MF417586 belong to different ‘sub-groups’ was therefore unclear, given that similar numbers of nucleotide substitutions between sequences (e.g., MF417602 and MF417587: SSA1-SG2; two transitions (i.e., 2xC/T), 0.37% nucleotide difference), and sequences with greater numbers of nucleotide substitutions (e.g., MF417590, MF417582: SSA1-SG1; three transitions (2xG/A, 1xT/C) and one transversion (A/T); 0.75% nucleotide difference) were placed within the same subgroups.

### Species delimitation and introgression

The genome-wide SNP-based Principal Component Analysis (PCA) discriminated between SSA1 and SSA2, suggesting that they are genetically distinct and likely to be different species, whereas samples from SSA4 clustered with SSA2 (Fig. 1). Three groupings were identified within SSA1, suggesting a level of intra-species substructure. These observations were further explored using ML phylogenies generated with RAxML. An initial such analysis based on the mtCOI sequences (Fig. 2A) broadly divided these 63 SSA individuals into three clusters containing the populations of (Wosula, et al. 2017): (i) SSA2 cluster containing individuals previously identified as either SSA4 or SSA2 by mtCOI gene sequences with 100% bootstrap confidence, (ii) SSA1 cluster of the populations defined as ‘SSA-CA/SSA-ESA’ (node confidence 89%), and (iii) SSA1 cluster that defined as ‘SSA-ECA/SSA-WA’ populations (node confidence 96%).

**Fig. 1.**
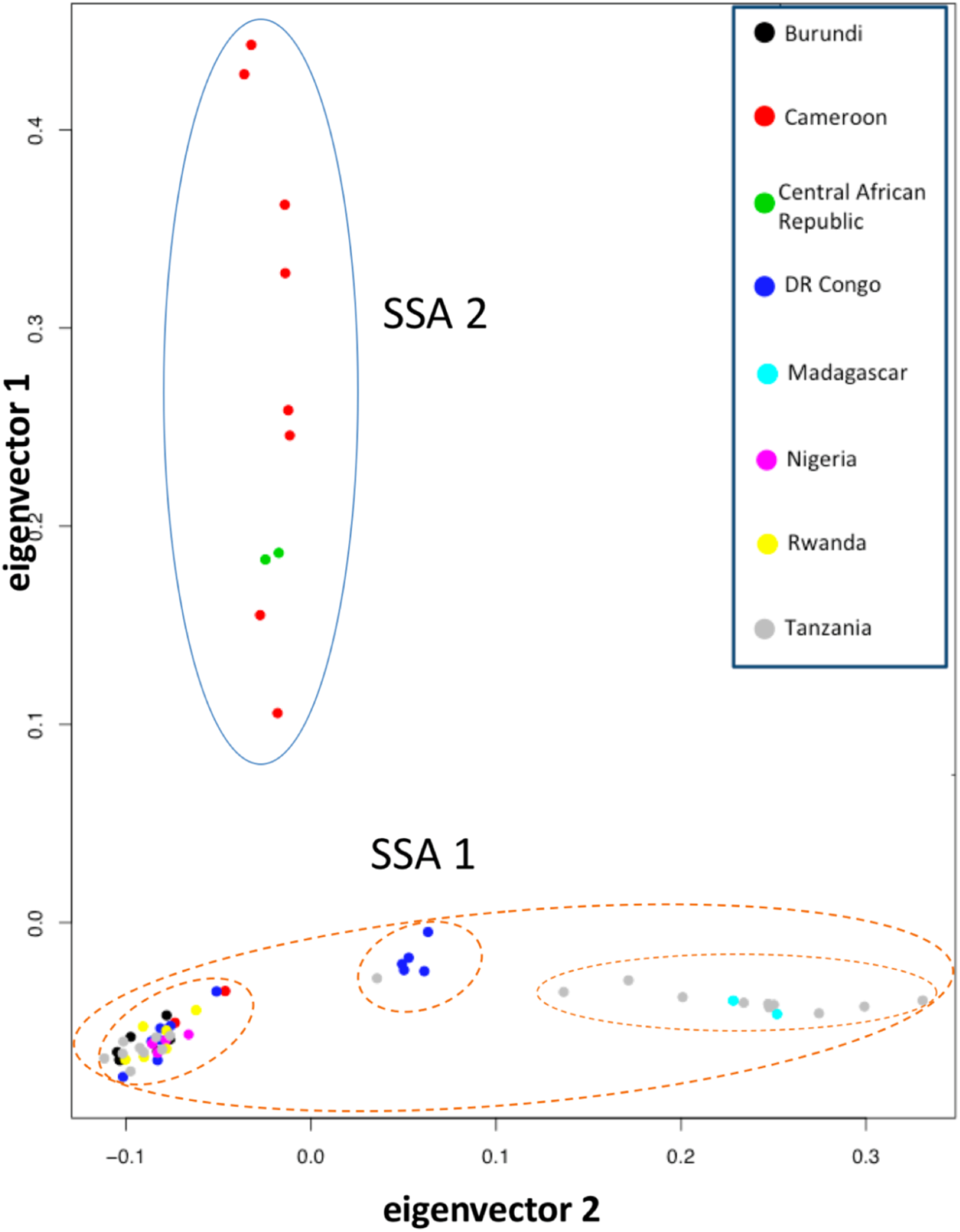
Principal component analysis of SSA *Bemisia* samples. Principal component analysis (PCA) was based on 14,358 genome-wide SNPs across the 63 sub-Saharan African cassava *Bemisia* samples. Clustering of SSA2 (blue circle) from SSA1 (red circles) supported their genetic distinctness as separate species. Three groupings are identified within SSA1, suggesting a level of intra-species substructure including the potential gene flow contact zone. Country of origins for these SSA1 and SSA2 *Bemisia* species are shown by unique colour codes.

**Fig. 2.**
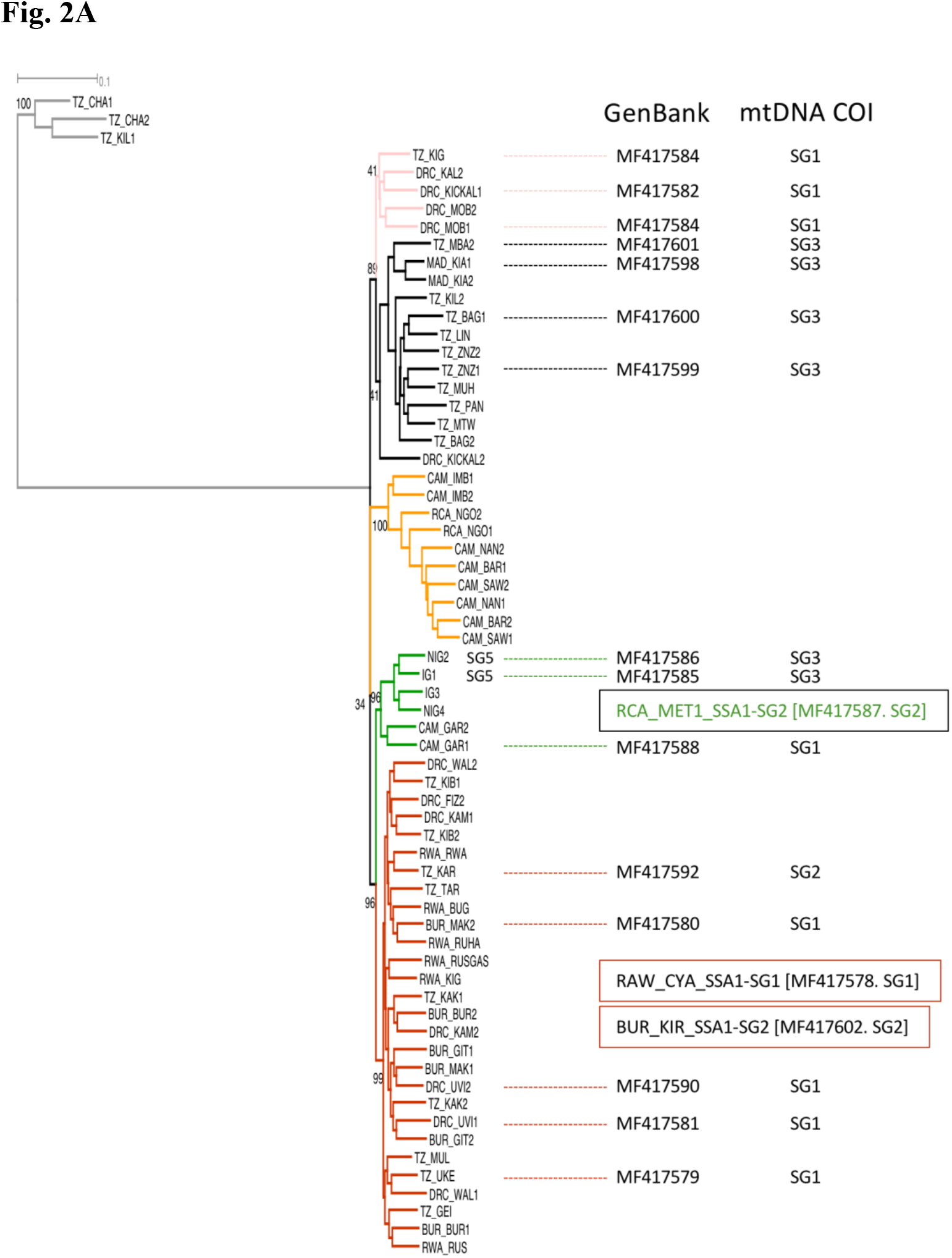
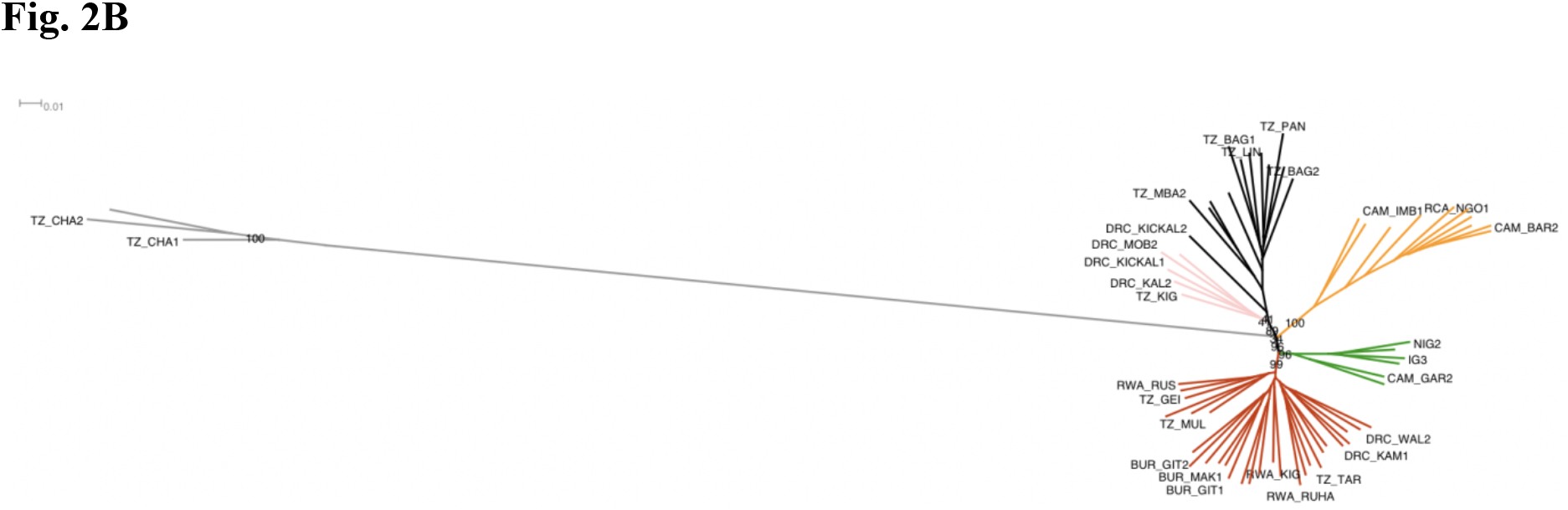
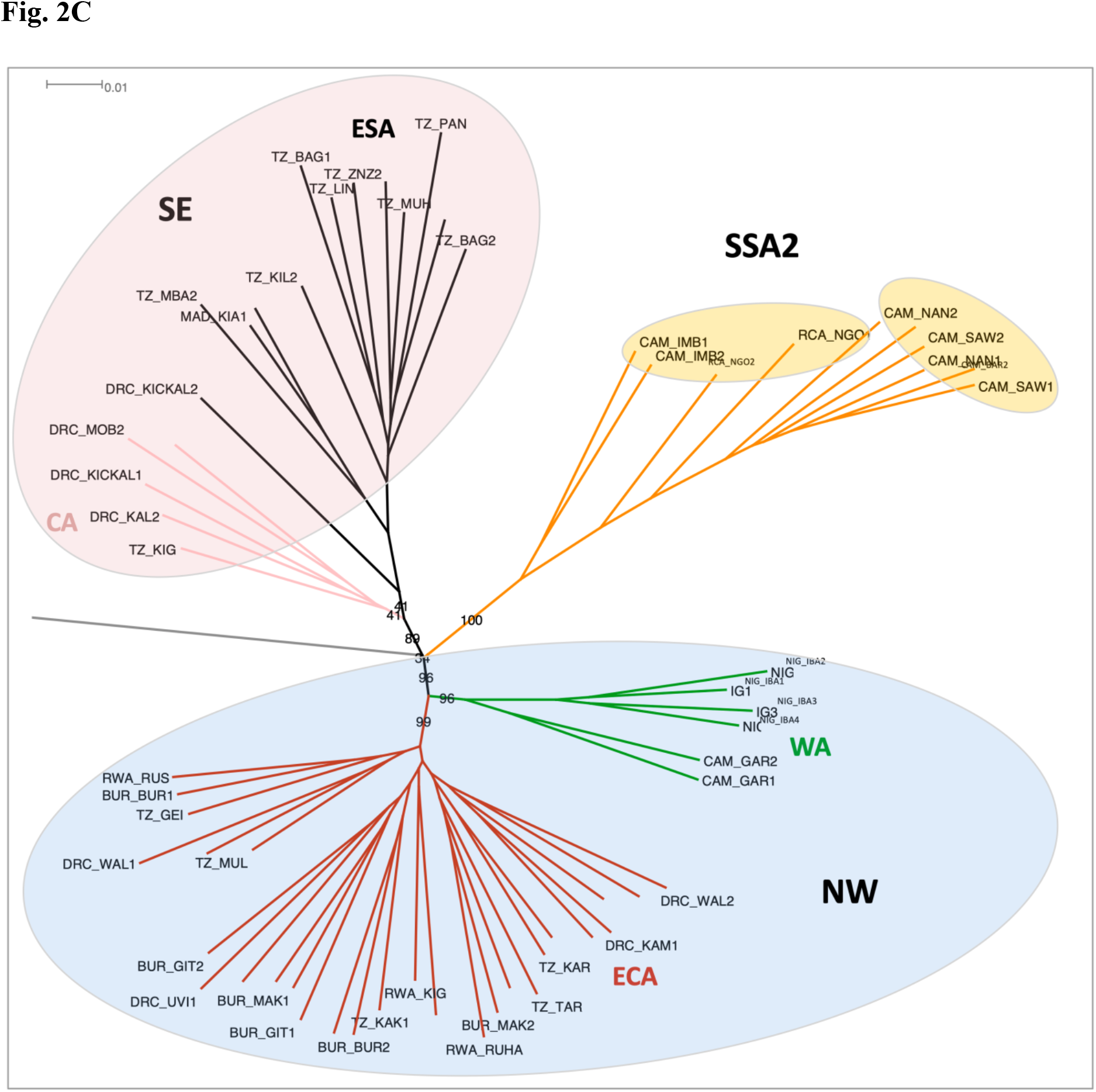
Phylogeny of SSA populations. **A:** Phylogeny based on mtCOI of 63 SSA *Bemisia* individuals. The phylogeny was constructed using RAxML with 1,000 bootstrap replications. Sample names are as in Fig. S1, and GenBank accession numbers for the sequences used, and corresponding mtCOI SSA1-subgroups (SG) of either SG1, SG2 and SG3, are listed on the right. The *Bemisia* SSA2-SSA4 clade is indicated by yellow branches, and the lineages corresponding to the SSA1 populations analysed are indicated as follows: CA (pink), ESA (black), WA (green) and ECA (red). **B:** Phylogeny based on 14,358 genome-wide SNPs of the individuals analysed in A. The phylogeny was constructed using RAxML with 1,000 bootstrap replications. Sample names and lineage colours are as for **A** above. **C:** Detail of the RAxML phylogeny in **B** showing only the SSA1 and SSA2-SSA4 lineages, with population names added. Bootstrap confidence percentages are visible for the main branches.

A comprehensive RAxML phylogenomic analysis based on the 14,358 genome-wide SNPs (Figs. 2B, 2C) confirmed SSA1 and SSA2 species status and is consistent with that earlier identified using partial mtCOI gene sequence identity. This analysis clearly showed SSA1 to comprise two clusters, denoted SSA1-SE (South-Eastern) and SSA1-NW (North-Western) (Fig. 2C), respectively, reflecting the geographic distribution of the populations identified within them (SSA1-SE comprising samples from Democratic Republic of Congo (DRC), Tanzania (TZ), and Madagascar (MAD); and SSA1-NW comprising samples from Cameroon (CMR), Nigeria (NGA), Democratic Republic of Congo (DRC), Rwanda (RWA), Burundi (BUR) and Tanzania (TZ)). Within SSA1-NW, the ‘SSA-ECA’ and ‘SSA-WA’ populations as defined by Wosula et al. (2017) are clustered separately with 99% and 96% confidence values. Within SSA1-SE, the ‘SSA-ESA’ and ‘SSA-CA’ populations as defined by Wosula et al. (2017) are partially resolved, sharing only 34% confidence and suggesting insufficient genomic evidence for this phylogenetic analysis to confirm them as separate genetic groups.

Admixture analysis identified three genetic clusters (Fig. 3) that separated the SSA2 species (highlighted in green) from the SSA1 species, which was again divided into two distinct genetic backgrounds corresponding to SSA1-NW (containing SSA-WA and SSA-ECA (highlighted in blue)) and SSA1-SE (containing SSA-CA and SSA-ESA (highlighted in red)). Some SSA2 and all the SSA1-CA samples showed evidence for low levels of admixture. Interestingly, *F*_st_ analysis (Table 1) supported the status of the two sub-groups SSA1-NW and SSA-SE as intermediate at the species-level differentiation between any of the SSA1 populations and the pooled SSA2 populations (the latter ranging from 0.28582 – 0.33406). This was particularly evident for SSA1-NW versus the SSA-ESA population of SSA1-SE, with *F*_st_ of 0.22605 - 0.24304. *F*_st_ differentiation was less clear for the more weakly defined SSA-CA population, whose *F*_st_ scores versus SSA1-NW were not significantly different to those between populations within the two major SSA1 groups. While the overall differentiation between SSA1-NW and SSA1-SE is consistent with them being sub-species of SSA1, the observations concerning the SSA-CA population suggest gene flow between CA and neighbouring populations from both sub-species.

**Fig. 3.**
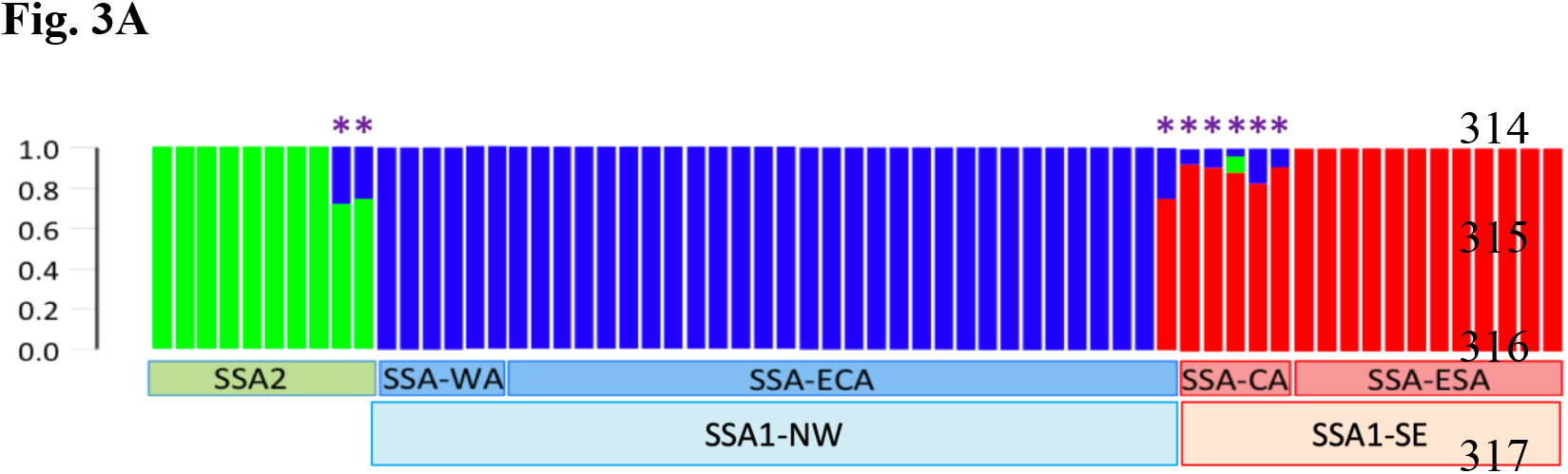
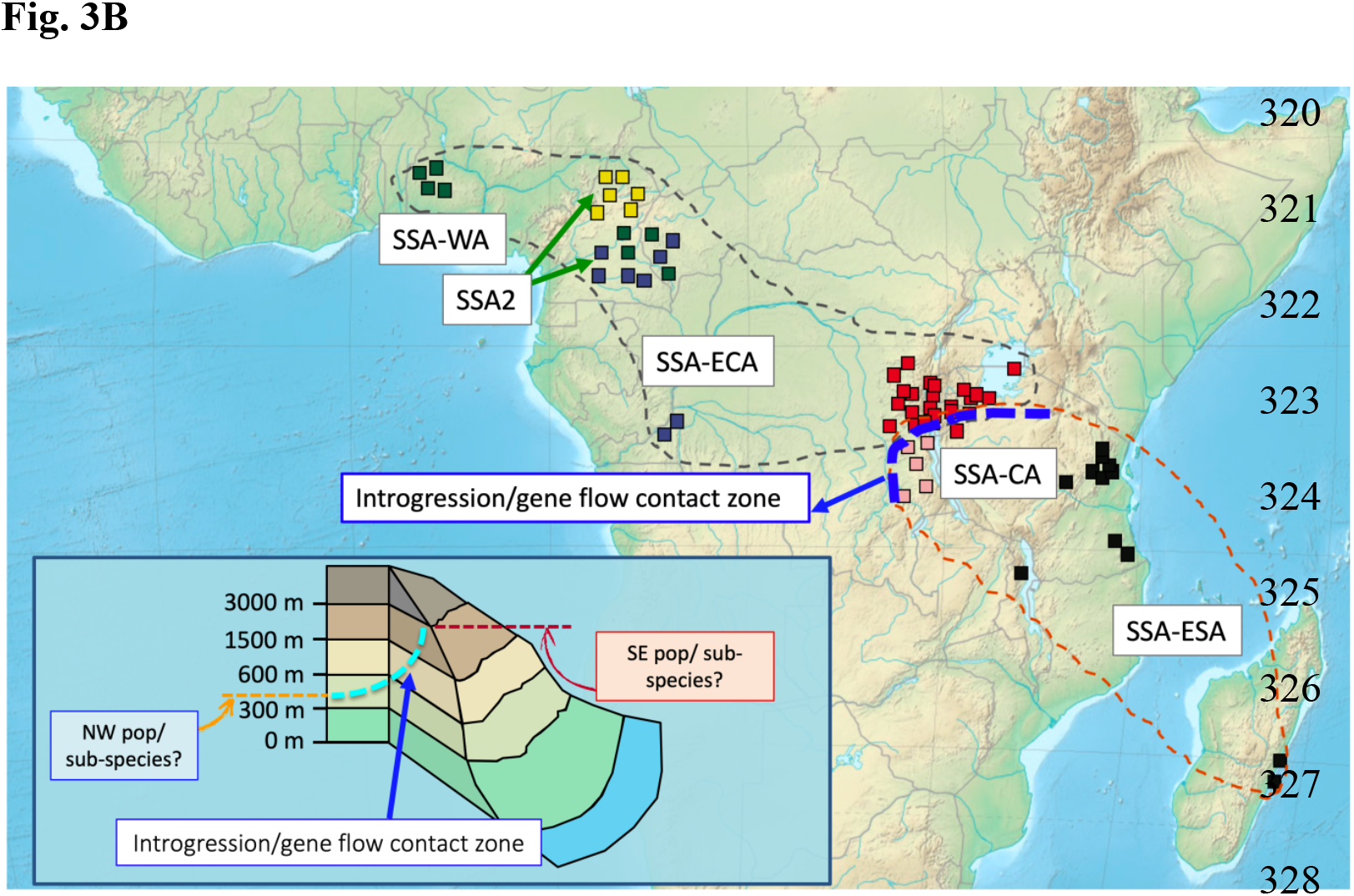
Populations and species of *Bemisia* across sub-Saharan Africa. Population structure analysis using Admixture of 63 individuals from the sub-Saharan *Bemisia* cryptic species clade based on 14,358 SNPs (K=3). Individuals were clustered into the SSA2 species group, as well as two genetic clusters (red, blue) that belonged to SSA1 based on partial mtCOI gene signature. The samples are shown on the map: green squares represent SSA-WA, red squares represent SSA-ECA, pink squares represent SSA-CA and black squares represent SSA-ECA individuals. Yellow and blue squares represent SSA2 and ‘SSA4’ individuals, respectively (the latter being NUMT artefacts of SSA2 – see text). One SSA1 genetic cluster in the Admixture panel included the SSA-WA and SSA-ECA groups (35 blue individuals and one red-blue individual), and the other SSA1 genetic cluster included five SSA-CA (blue-red/blue-red-green) and 12 SSA-ESA (red) individuals. Evidence of gene flow between SSA1 genetic clusters were predominantly detected in SSA-CA individuals (four red-blue individuals in the Admixture analysis). Three inter-specific hybrids (two from SSA2; one from SSA-CA with SSA1 (red-blue) and SSA2 (green) genetic backgrounds) were also detected in the SSA *Bemisia* individuals analysed. The gene flow contact zone between populations on the NW-SE transect is indicated by the thick blue dashed line. SSA1-NW and SSA1-SE sub-species boundary lines are indicative only. Landscape elevations across the sampling sites are shown in insert.

Within sub-species SSA1-NW, the SSA-WA population is predominantly found in the lowlands (*ca*. 300m - 1500m), whereas the SSA-ECA/SSA-CA populations were found at higher altitudes (from *ca*. 600m - ≥1500m). Individuals belonging to the SSA-ESA group were predominantly from coastal landscape of Tanzania and Madagascar (Fig. 3). A gene flow contact zone was detected at the SSA-CA region represented by individuals from DRC (n = 5), and one TZA individual from SSA-ECA group. Hybridization was detected in two SSA2 individuals (green/blue genetic backgrounds) and in one DRC individual from SSA-CA (red/green/blue in Fig. 3).

Results from our D-statistics test (ABBA-BABA test) between three groups [ESA+CA], [SSA2+SSA4] and [ECA+WA] using MEAM1 as an outgroup identified strong signals of introgression between the [ESA+CA] and [ECA+WA] groups which belonged to the same SSA1 species based on mtCOI gene analysis, where the strong introgression signals between [ESA+CA] and [ECA+WA] could potentially represent well-defined sub-populations/sub-species occupying different geographical landscapes, with gene flow occurring between these sub-populations/sub-species at the contact zone (Fig. 4).

**Fig. 4.**
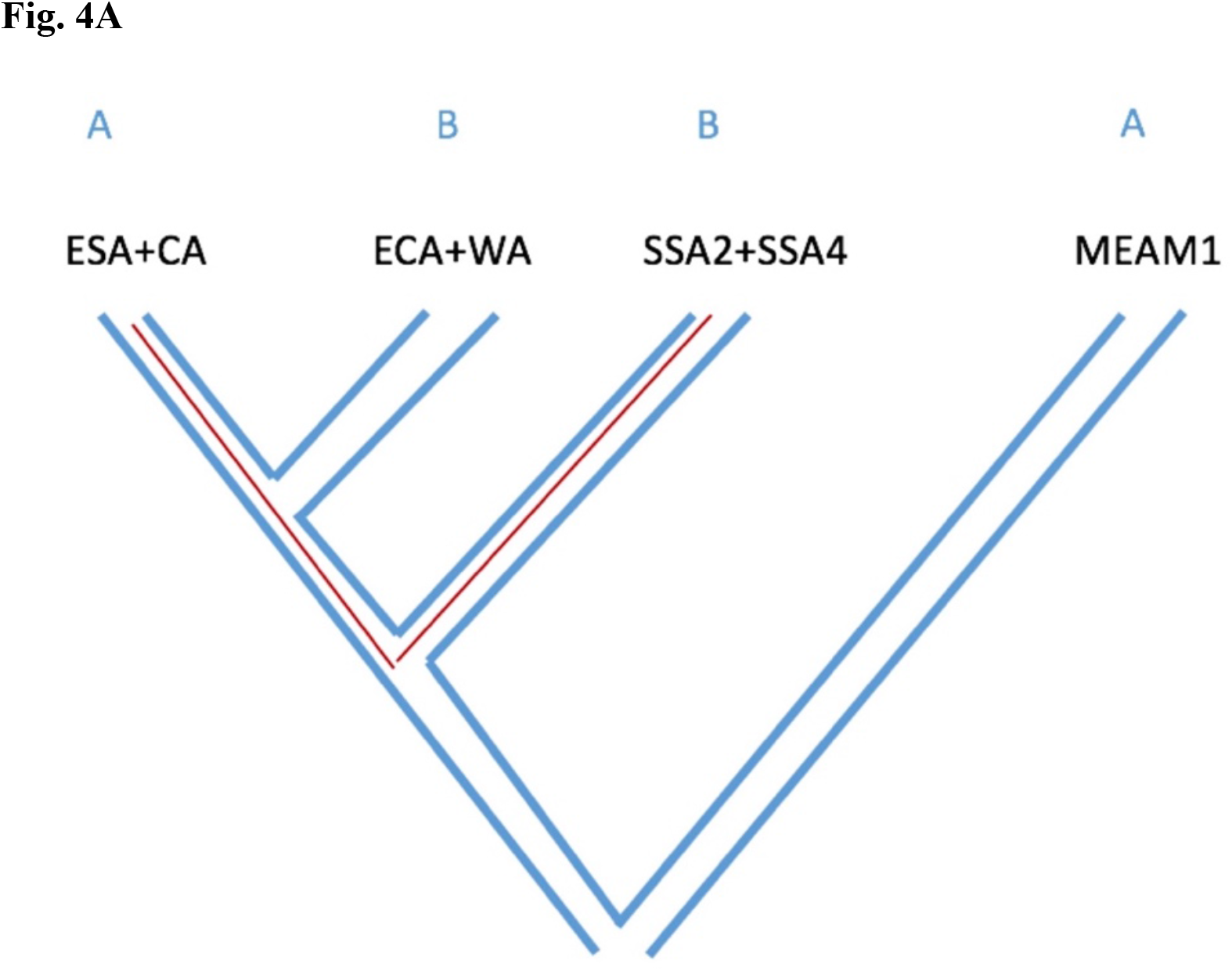
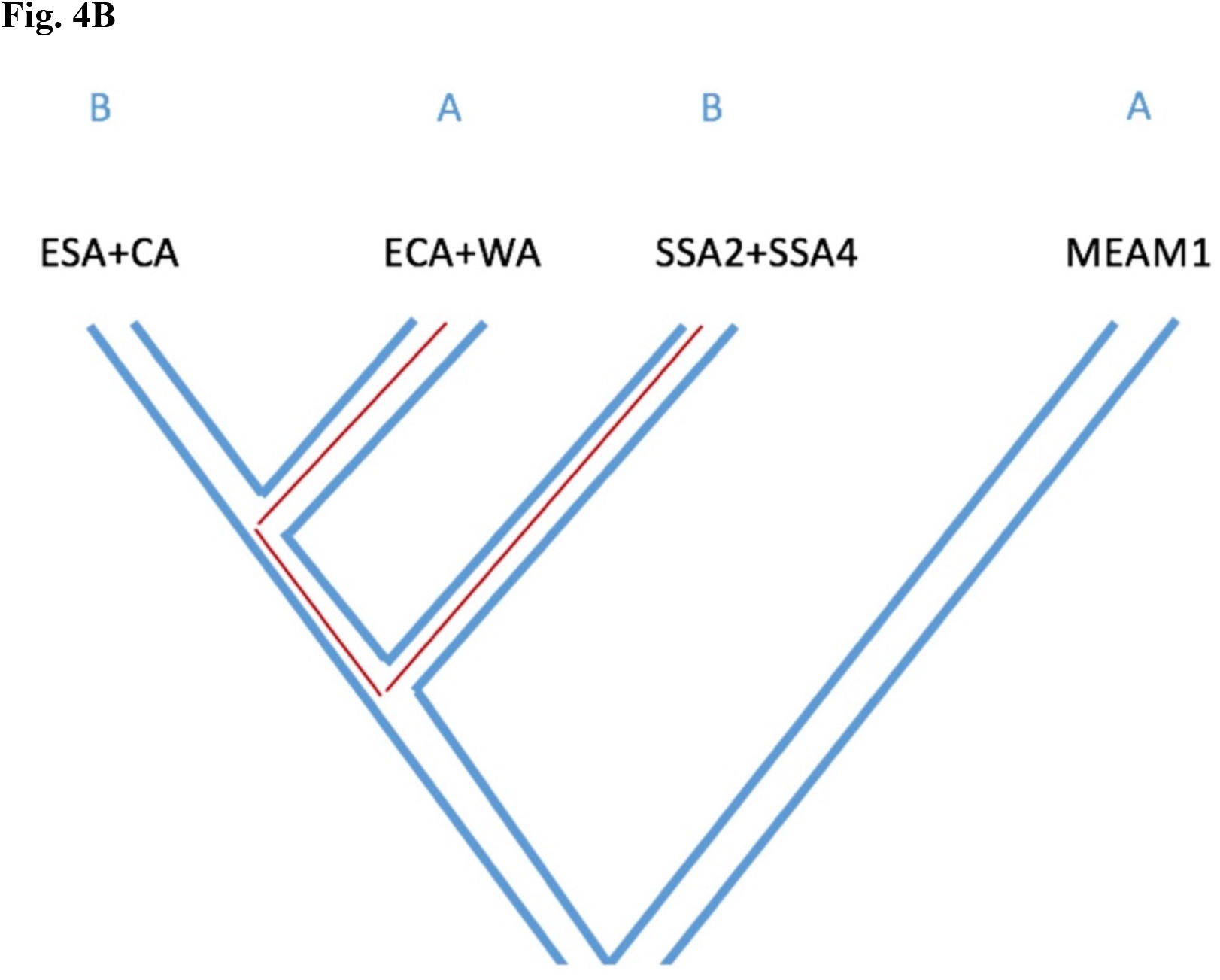
Introgression site patterns using a four-Taxon ABBA/BABA test. **A.** The ABBA and BABA sites used in the test are derived from ESA+CA and ECA+WA belonging to SSA1 species, compared to the SSA2+ SSA4 (**Fig. 4B**) group which are previously reported as two distinct species but were identified as belonging to one species (SSA2) in this study.

The test also detects signals of introgression within the same geographical region between the SSA2 and SSA1 (i.e., [ECA+WA]) species. The same signatures of interspecies admixture were also evident from the Treemix results (Fig. 5). The ML tree generated in Treemix also supported the SSA4 species as part of SSA2. The populations/subspecies SSA-ECA and SSA-WA cluster together which is consistent with their geographical origin (i.e., SSA1-NW), whereas the SSA-CA population clusters with the SSA-ECA population (i.e., representing the south-eastern geographic origin). Gene flow between these SSA1 populations is especially evident at the contact zone (i.e., at the SSA-CA geographical location). The migration edges (m = 2) revealed gene flow between SSA2 and the SSA1 species from central Africa (i.e., SSA-CA) and supports detection of hybrid individuals identified in the admixture analysis (Fig. 3).

**Fig. 5.**
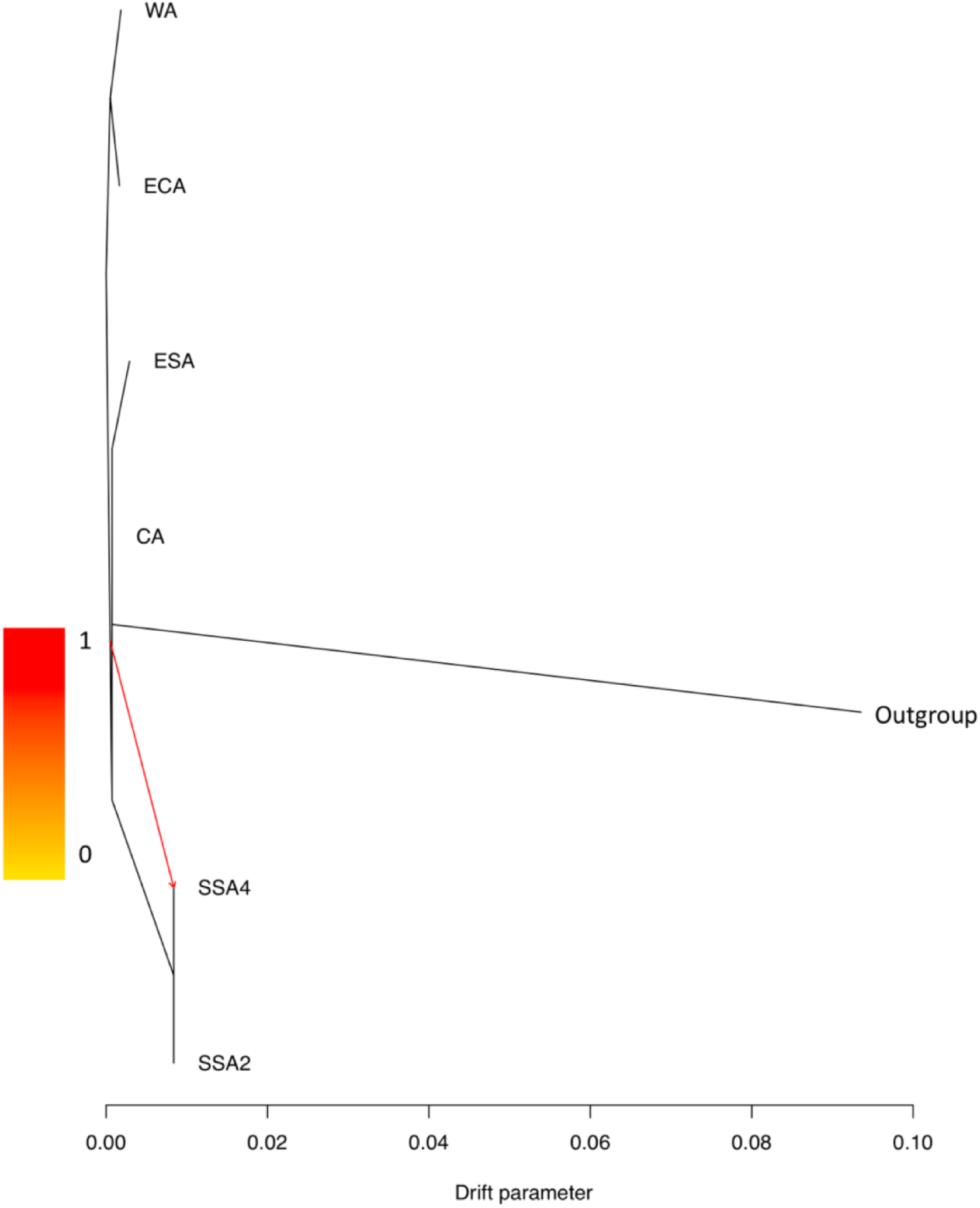
Gene flow between SSA species. TreeMix analysis showing gene flow from SSA1 to SSA2/SSA4 (red arrow). Clustering between ECA and WA, and between ESA and CA indicated close evolutionary relationship. SSA2 and SSA4 shared a genetic clade with no branch depth indicating these two species were identical.

## Discussion

We combined genome-wide SNPs and mtCOI markers to gain unprecedented evolutionary ecological and population genomic insights into the sub-Saharan African cassava whitefly species complex. By taking into consideration the impact of NUMTs, we showed that there is no conflict between nuclear and mitochondrial DNA markers in terms of defining species status, contrary to the study of Wosula et al. (2017) based on the same data set. A thorough filtering of genome-wide SNP markers also discerned the evolutionary forces shaping incipient speciation and admixture in the *Bemisia* SSA1 whitefly species. Various evolutionary genetic studies based on the partial mtCOI gene have proposed the presence of population subgroups (e.g., Ghosh, et al. 2018; Legg, et al. 2014) in the SSA1 species. Based on genome-wide SNP markers, we found no evidence to support the subgroup classification, although for the first time, the SSA1 species as a whole, could be broadly divided into two well-defined north western (NW) and south eastern (SE) populations/sub-species, with a contact zone separated by the eastern highland and central/western lowland regions. Evidence for interspecies hybridisation was also detected between the SSA1 and SSA2 species, where bidirectional genetic contributions were observed between SSA2 and SSA1, and will have important implications to understanding the evolutionary genetics associated between plant virus transmission capacity and the socioeconomically important SSA *Bemisia* whitefly pest complex.

In the study of Elfekih et al. (2018a), mtCOI sequences from the MEAM1, MED, Indian Ocean (IO) and Australia (AUS) *B. tabaci* cryptic species were interrogated prior to generating nextRAD genome-wide SNP data to investigate the history of global introduction pathways of the MED and MEAM1 species. Elfekih et al. (2018a), Elfekih et al. (2015), Elfekih et al. (2016a), and Elfekih et al. (2016b) showed that species status of these cryptic *B. tabaci* species as ascertained by the partial mtCOI gene was corroborated by genome-wide SNP data. Similar to this study, genome-wide SNP markers also detected genomic signatures of hybridization between the MED, MEAM1 and IO, while also revealing the role played by NUMTs in confounding our effort to define species status such as between the MEAM1 and MEAM2 species (Delatte, et al. 2005; Elfekih, et al. 2018a; Tay, et al. 2017a).

Wosula et al. (2017) adopted the nextRAD genome-wide sequencing method outlined by Elfekih et al. (2015), Elfekih et al. (2016a), and Elfekih et al. (2016b) to investigate gene flow patterns and diversity in the African cassava whitefly *Bemisia* SSA1, SSA2, SSA4 species complex, but overlooked the impact of NUMTs associated with the *B. tabaci* mtCOI partial sequences. This oversight led to the major conclusion that the widely used mtCOI gene was an unsuitable marker for differentiating between African *Bemisia* cryptic species. Increasingly, reanalyses of the *B. tabaci* complex cryptic species partial mtCOI gene dataset have detected the presence of significant NUMTs and pseudogenes (Kunz, et al. 2019b; Tay, et al. 2017a; Vyskočilová, et al. 2018). These studies highlighted the challenges faced by researchers to delimit species in the cryptic *B. tabaci* species complex, despite the promises of molecular systematics and species delimitation via mitochondrial barcoding approaches. Similar to MEAM2 (Boykin, et al. 2013; Dinsdale, et al. 2010), the *Bemisia* cryptic species SSA4 (Berry, et al. 2004) was shown to be the product of NUMTs. This species misidentification underpinned all subsequent interpretations of gene flow and evolutionary ecology in the SSA *Bemisia* cassava whitefly species (Wosula, et al. 2017) based on genome-wide SNP markers.

Significant genetic diversity in the *B. tabaci* species complex can obscure species and sub-species identification. For instance, Elfekih et al. (2018a) detected significant diversity in the MED species based on nextRAD SNP data; however, the MED-ASL species was not identified until confirmed by complete mitogenome study, mating experiments (Vyskočilová, et al. 2018), and host preference studies (Vyskočilová, et al. 2019). Similar studies are needed to provide additional biological support for subgroup/sub-species status within SSA1.

In developing the DNA barcoding approach, Hebert et al. (2003) showed that the COI gene is especially effective at differentiating at the species, order and phylum levels due to the associated nucleotide substitutional rates for this gene, while for fine-scale identification such as within species level, the use of genes with greater substitutional rates were recommended. Subgroup classification within the SSA1 African cassava whitefly species based on the mtDNA gene might be possible, provided that a gene region with an appropriate substitutional rate is identified. Nevertheless, there is currently no evidence for maternal lineage-based patterns to population dispersal behaviour, host adaptation, and fitness cost differentiation in the *Bemisia tabaci* complex and the African cassava *Bemisia* species complex, and therefore the subgroup delimitation based on the partial mtCOI gene is not justified.

Previously, Legg et al. (2014) reported that the SSA1-SG1, representing the major SSA1 mtCOI subgroup, was implicated in the severe CMD pandemic in East Africa. Based on the subgroup classification, Wosula et al. (2017) reclassified these SSA1-SG1 individuals into three SNP-based groups of SSA-ECA, SSA-CA, and SSA-WA due to perceived inconsistencies (e.g., SSA1-SG3, SSA1-SG5; see Ghosh et al. (2015)) based solely on the utilization of the short (i.e., *ca*. 657bp) mtCOI partial gene marker. Based on the same dataset, our reanalysis found that while phylogenetic signals were detected for the separate clustering of the ‘WA’ and ‘ECA’ populations, there was not a strong genetic difference between the ‘CA’ and ‘ESA’ populations, and we therefore caution against the adoption of the nomenclature as proposed by Wosula et al. (2017) in future studies. Furthermore, admixture analysis also showed a substantial gene flow limitation between the ‘WA/ECA’ groups and the ‘CA/ESA’ groups.

Although adopting the subgroup status could help resolve virus transmission capacity and provide insights into understanding pest introduction pathways (Anderson, et al. 2018; Tabachnick 1996), there is currently insufficient evidence to link either the SSA1-NW (i.e., ‘WA/ECA’) or SSA1-SW (i.e., ‘CA/ESA’) populations/subgroups to historical CMD outbreaks. Legg et al. (2002) described a distinct genetic cluster (*B. tabaci* Ug2) assumed to be associated with the CMD epidemic in the 1990’s, mainly due to its geographic distribution in relation to the front of the epidemic. However, transmission leading to outbreaks of CMD in the sub-Saharan Africa region is complex, with one factor being the sharing and propagation of diseased cassava cuttings between farmers. There was a recombination event between two geminiviruses that produced a more virulent hybrid (Colvin, et al. 2004), however we have little historical information about how the viruses interacted with the different *Bemisia* cryptic species in terms of disease dynamics (Polston, et al. 2014). The role of the SSA2 species in the CMD epidemic is also unclear given that the SSA2’s habitat range (east and west Africa) overlaps the SSA1 species (Legg, et al. 2002; Mugerwa, et al. 2018; Wosula, et al. 2017). The exact relationship between SSA1/SSA2 and outbreaks of CMD (Manani, et al. 2017) should be re-investigated, taking into consideration the current nextRAD SNP data as genomic evidence that supports the presence of hybrid SSA1/SSA2 individuals. Furthermore, the vector competencies of other common cassava whitefly species (such as *Bemisia afer*) to different cassava viruses also needs to be fully understood.

Stringent delimitation of population groups across the full habitat range of the SSA1 species will be needed to ascertain whether the SSA1 as a whole is represented by genetically distinct groups where limited gene flow could be further identified to external (e.g., geographic, climatic) and/or metagenomic (e.g., secondary symbionts) factors. The genetic differentiation between the two SSA1 genetic groups suggests they can be considered as subspecies of SSA1. Recognition of such biological classifications and their distribution in sub-Saharan Africa will be important for management of these agriculturally and biosecurity significant species, as for others posing a threat to global health and biosecurity (Paini, et al. 2016).

The South-Eastern SSA1 sub-populations belonging to Madagascar (SSA-ESA) and Tanzania (SSA-CA) are geographically isolated from the North-Western sub-populations sampled from Nigeria (SSA-WA), Democratic Republic of Congo and Burundi (SSA-ECA). The mountain range separating these two SSA1 categories (SE and NW sub-populations) poses a geographical barrier to gene flow, and representing a potential factor that could promote incipient allopatric speciation. For instance, the East African Rift Valley is a major geographical barrier providing the right environment for incipient species and recent radiations to thrive and evolve (Gottelli, et al. 2004). Specifically, the western branch of the Rift Valley, i.e., the Albertine Rift covering part of Tanzania, Burundi, Rwanda, Uganda and Democratic Republic of Congo, is a chain of mountains situated at 1500–3500 meters above sea level, and formed by volcanic activity that started during the plio-pleistocene. In addition to the volcanic activity, climate change during this glacial period led to habitat fragmentation and therefore species diversification in the newly formed biogeographical zones (Huhndorf, et al. 2007). In addition to allopatric forces, species diversification in the whitefly cryptic species system has been explained by host plant associations and the ability of this system to feed on a wide range of host plants including acquiring new ones (Matsubayashi, et al. 2010). Using RNAseq, Malka et al. (2018) provided insights into how the variation in level of expression of detoxification genes might have conferred plasticity to the complex and that likely also contributed to its diversification.

*B. tabaci sensu lato* on cassava in sub-Saharan Africa is a species complex going through a continuous diversification process. The examination of the genetic diversity via mitochondrial and genome-wide markers validated discrimination between SSA1 and SSA2, and the latter being the same as SSA4. This was mainly facilitated by careful data analysis, and taking into consideration artefacts caused by pseudogenes (NUMTs). The SSA1 species shows signatures of incipient speciation with sub-populations divided into two biogeographical niches; North Western (NW) and South Eastern (SE) regions with a hybrid zone between Tanzania, Uganda and Democratic Republic of Congo. We conclude that accurate species identification that takes into consideration all the evolutionary forces driving the diversification in this system will be critical in the implementation of any future management strategy for this invasive and economically important insect pest complex. Further comparative analyses on the endosymbiont metacommunities and host association can shed more light onto the major causes underlying the gradual process of species diversification and adaptive radiations in the cassava whitefly system.

## Acknowledgements

SE was funded by the CSIRO Office of the Chief Executive (OCE) post-doctoral fellowship (R-4546-1) and a European Molecular Biology Organization (EMBO) fellowship ASTF-6889. WTT, SE, KG DK and TW were supported by CSIRO H&B Genes of Biosecurity Importance fund (R-8681-1). SM, AP, WTT, SV, JC, PDB were supported by the Natural Resources Institute, University of Greenwich from a grant provided by the Bill & Melinda Gates Foundation (Grant Agreement OPP1058938).

## Literature cited

Alexander DH, Novembre J, Lange K 2009. Fast model-based estimation of ancestry in unrelated individuals. Genome Research 19: 1655–1664. doi: 10.1101/gr.094052.109

Anderson CJ, et al. 2018. Hybridization and gene flow in the mega-pest lineage of moth, *Helicoverpa*. Proc Natl Acad Sci U S A 115: 5034–5039. doi: 10.1073/pnas.1718831115

Anderson CJ, Tay WT, McGaughran A, Gordon K, Walsh TK 2016. Population structure and gene flow in the global pest, *Helicoverpa armigera*. Mol Ecol 25: 5296–5311. doi: 10.1111/mec.13841

Behere GT, Tay WT, Russell DA, Batterham P 2008. Molecular markers to discriminate among four pest species of *Helicoverpa* (Lepidoptera: Noctuidae). Bull Entomol Res 98: 599–603. doi: 10.1017/S0007485308005956

Berry SD, et al. 2004. Molecular evidence for five distinct *Bemisia tabaci* (Homoptera: Aleyrodidae) geographic haplotypes associated with cassava plants in sub-Saharan Africa. Annals of the Entomological Society of America 97: 852–859. doi: Doi 10.1603/0013-8746(2004)097[0852:Meffdb]2.0.Co;2

Bolger AM, Lohse M, Usadel B 2014. Trimmomatic: a flexible trimmer for Illumina sequence data. Bioinformatics 30: 2114–2120. doi: 10.1093/bioinformatics/btu170

Boykin LM, Bell CD, Evans G, Small I, De Barro PJ 2013. Is agriculture driving the diversification of the *Bemisia tabaci* species complex (Hemiptera: Sternorrhyncha: Aleyrodidae)?: Dating, diversification and biogeographic evidence revealed. BMC Evol Biol 13: 228. doi: 10.1186/1471-2148-13-228

Boykin LM, Savill A, De Barro P 2017. Updated mtCOI reference dataset for the Bemisia tabaci species complex F1000Res 6: 1835. doi: 10.12688/f1000research.12858.1

Chen WB, et al. 2016. The draft genome of whitefly *Bemisia tabaci* MEAM1, a global crop pest, provides novel insights into virus transmission, host adaptation, and insecticide resistance. BMC Biology 14: 110. doi: 10.1186/s12915-016-0321-y

Colvin J, Omongo CA, Maruthi MN, Otim-Nape GW, Thresh JM 2004. Dual begomovirus infections and high *Bemisia tabaci* populations: two factors driving the spread of a cassava mosaic disease pandemic. Plant Pathology 53: 577–584. doi: 10.1111/j.1365-3059.2004.01062.x

De Barro PJ, Liu SS, Boykin LM, Dinsdale AB 2011. *Bemisia tabaci*: a statement of species status. Annu Rev Entomol 56: 1–19. doi: 10.1146/annurev-ento-112408-085504

Delatte H, et al. 2005. A new silverleaf-inducing biotype Ms of *Bemisia tabaci* (Hemiptera: Aleyrodidae) indigenous to the islands of the south-west Indian Ocean. Bulletin of Entomological Research 95: 29–35. doi: 10.1079/Ber2004337

Dinsdale A, Cook L, Riginos C, Buckley YM, De Barro P 2010. Refined Global Analysis of *Bemisia tabaci* (Hemiptera: Sternorrhyncha: Aleyrodoidea: Aleyrodidae) Mitochondrial Cytochrome Oxidase 1 to Identify Species Level Genetic Boundaries. Annals of the Entomological Society of America 103: 196–208. doi: 10.1603/An09061

Eaton DAR 2014. PyRAD: assembly of de novo RADseq loci for phylogenetic analyses. Bioinformatics 30: 1844–1849. doi: 10.1093/bioinformatics/btu121

Elfekih S, et al. 2018a. Genome-wide analyses of the *Bemisia tabaci* species complex reveal contrasting patterns of admixture and complex demographic histories. PLoS One 13: e0190555. doi: 10.1371/journal.pone.0190555

Elfekih S, et al. 2015. Evolutionary genomics of *Bemisia tabaci* and characterization of its endosymbiont metacommunities using nextRAD sequencing. International Plant and Animal Genome Asia 2015 July 12-15th, 2015: Singapore.

Elfekih S, et al. 2016a. Genome-Wide SNPs Decipher Global Incursion Pathways in the *Bemisia tabaci* Species Complex. International Plant and Animal Genome Conferences. 2016a 9-13 January 2016: San Diego, CA, U. S. A.

Elfekih S, et al. 2016b. Genome-wide scans unravel fine-scale invasion routes in the *Bemisia tabaci* species complex. 2nd International Whitefly Symposium,. 2016b 14-19 February 2016 Arusha, Tanzania. p38.

Elfekih S, Tay WT, Gordon K, Court LN, De Barro PJ 2018b. Standardized molecular diagnostic tool for the identification of cryptic species within the *Bemisia tabaci* complex. Pest Manag Sci 74: 170–173. doi: 10.1002/ps.4676

FAOSTAT 2017 [cited 2018 Jan-2018]. Available from: http://www.fao.org/faostat/en/-data/QC/visualize

Fumagalli M, et al. 2013. Quantifying Population Genetic Differentiation from Next-Generation Sequencing Data. Genetics 195: 979-+. doi: 10.1534/genetics.113.154740

Ghosh S, Bouvaine S, Maruthi MN 2015. Prevalence and genetic diversity of endosymbiotic bacteria infecting cassava whiteflies in Africa. BMC Microbiology 15: 93. doi: 10.1186/s12866-015-0425-5

Ghosh S, Bouvaine S, Richardson SCW, Ghanim M, Maruthi MN 2018. Fitness costs associated with infections of secondary endosymbionts in the cassava whitefly species *Bemisia tabaci*. Journal of Pest Science 91: 17–28. doi: 10.1007/s10340-017-0910-8

Gottelli D, Marino J, Sillero-Zubiri C, Funk SM 2004. The effect of the last glacial age on speciation and population genetic structure of the endangered Ethiopian wolf (*Canis simensis*). Molecular Ecology 13: 2275–2286. doi: 10.1111/j.1365-294X.2004.02226.x

Hadjistylli M, Roderick GK, Gauthier N 2015. First report of the Sub-Saharan Africa 2 species of the *Bemisia tabaci* complex in the Southern France. Phytoparasitica 43: 679–687. doi: 10.1007/s12600-015-0480-3

Hebert PDN, Cywinska A, Ball SL, DeWaard JR 2003. Biological identifications through DNA barcodes. Proceedings of the Royal Society B-Biological Sciences 270: 313–321. doi: 10.1098/rspb.2002.2218

Huhndorf MH, Peterhans JCK, Loew SS 2007. Comparative phylogeography of three endemic rodents from the Albertine Rift, east central Africa. Molecular Ecology 16: 663–674. doi: 10.1111/j.1365-294X.2007.03153.x

Katoh K, Standley DM 2013. MAFFT multiple sequence alignment software version 7: improvements in performance and usability. Mol Biol Evol 30: 772–780. doi: 10.1093/molbev/mst010

Korneliussen TS, Albrechtsen A, Nielsen R 2014. ANGSD: Analysis of Next Generation Sequencing Data. Bmc Bioinformatics 15: 356. doi: 10.1186/s12859-014-0356-4

Kunz D, et al. 2019a. Draft mitochondrial DNA genome of a 1920 Barbados cryptic *Bemisia tabaci* ‘New World’ species (Hemiptera: Aleyrodidae). Mitochondrial DNA Part B-Resources 4: 1183–1184. doi: 10.1080/23802359.2019.1591197

Kunz D, Tay WT, Elfekih S, Gordon KHJ, De Barro PJ 2019b. Take out the rubbish - Removing NUMTs and pseudogenes from the *Bemisia tabaci* cryptic species mtCOI database. Sci Rep Submitted.

Lee W, Park J, Lee GS, Lee S, Akimoto S 2013. Taxonomic status of the *Bemisia tabaci* complex (Hemiptera: Aleyrodidae) and reassessment of the number of its constituent species. PLoS One 8: e63817. doi: 10.1371/journal.pone.0063817

Legg JP, French R, Rogan D, Okao-Okuja G, Brown JK 2002. A distinct *Bemisia tabaci* (Gennadius) (Hemiptera: Sternorrhyncha: Aleyrodidae) genotype cluster is associated with the epidemic of severe cassava mosaic virus disease in Uganda. Molecular Ecology 11: 1219–1229. doi: DOI 10.1046/j.1365-294X.2002.01514.x

Legg JP, et al. 2014. Spatio-temporal patterns of genetic change amongst populations of cassava *Bemisia tabaci* whiteflies driving virus pandemics in East and Central Africa. Virus Res 186: 61–75. doi: 10.1016/j.virusres.2013.11.018

Li H, Durbin R 2010. Fast and accurate long-read alignment with Burrows-Wheeler transform. Bioinformatics 26: 589–595. doi: 10.1093/bioinformatics/btp698

Li H, et al. 2009. The Sequence Alignment/Map format and SAMtools. Bioinformatics 25: 2078–2079. doi: 10.1093/bioinformatics/btp352

Liu SS, et al. 2007. Asymmetric mating interactions drive widespread invasion and displacement in a whitefly. Science 318: 1769–1772. doi: 10.1126/science.1149887

Macfadyen S, et al. 2018. Cassava whitefly, *Bemisia tabaci* (Gennadius) (Hemiptera: Aleyrodidae) in East African farming landscapes: a review of the factors determining abundance. Bull Entomol Res 108: 565–582. doi: 10.1017/S0007485318000032

Malka O, et al. 2018. Species-complex diversification and host-plant associations in *Bemisia tabaci*: A plant-defence, detoxification perspective revealed by RNA-Seq analyses. Molecular Ecology 27: 4241–4256. doi: 10.1111/mec.14865

Manani DM, Ateka EM, Nyanjom SRG, Boykin LM 2017. Phylogenetic Relationships among Whiteflies in the *Bemisia tabaci* (Gennadius) Species Complex from Major Cassava Growing Areas in Kenya. Insects 8: 25. doi: 10.3390/insects8010025

Martin JH, Mound LA 2007. An annotated check list of the world’s whiteflies (Insecta: Hemiptera: Aleyrodidae). Zootaxa: 1–84.

Matsubayashi KW, Ohshima I, Nosil P 2010. Ecological speciation in phytophagous insects. Entomologia Experimentalis Et Applicata 134: 1–27. doi: 10.1111/j.1570-7458.2009.00916.x

Minato N, et al. 2019. Surveillance for Sri Lankan cassava mosaic virus (SLCMV) in Cambodia and Vietnam one year after its initial detection in a single plantation in 2015. PLoS One 14: e0212780. doi: 10.1371/journal.pone.0212780

Mugerwa H, et al. 2018. African ancestry of New World, *Bemisia tabaci*-whitefly species. Sci Rep 8: 2734. doi: 10.1038/s41598-018-20956-3

Paini DR, et al. 2016. Global threat to agriculture from invasive species. Proceedings of the National Academy of Sciences of the United States of America 113: 7575–7579. doi: 10.1073/pnas.1602205113

Patil BL, Fauquet CM 2009. Cassava mosaic geminiviruses: actual knowledge and perspectives. Mol Plant Pathol 10: 685–701. doi: 10.1111/j.1364-3703.2009.00559.x

Pickrell JK, Pritchard JK 2012. Inference of Population Splits and Mixtures from Genome-Wide Allele Frequency Data. PLoS Genetics 8: e1002967. doi: 10.1371/journal.pgen.1002967

Polston JE, De Barro P, Boykin LM 2014. Transmission specificities of plant viruses with the newly identified species of the *Bemisia tabaci* species complex. Pest Management Science 70: 1547–1552. doi: 10.1002/ps.3738

Reich D, Thangaraj K, Patterson N, Price AL, Singh L 2009. Reconstructing Indian population history. Nature 461: 489–U450. doi: 10.1038/nature08365

Stamatakis A 2006. RAxML-VI-HPC: Maximum likelihood-based phylogenetic analyses with thousands of taxa and mixed models. Bioinformatics 22: 2688–2690. doi: 10.1093/bioinformatics/btl446

Tabachnick WJ 1996. Culicoides variipennis and bluetongue-virus epidemiology in the United States. Annual Review of Entomology 41: 23–43. doi: DOI 10.1146/annurev.en.41.010196.000323

Tay WT, et al. 2017a. The Trouble with MEAM2: Implications of Pseudogenes on Species Delimitation in the Globally Invasive *Bemisia tabaci* (Hemiptera: Aleyrodidae) Cryptic Species Complex. Genome Biology and Evolution 9: 2732–2738. doi: 10.1093/gbe/evx173

Tay WT, et al. 2017b. Novel molecular approach to define pest species status and tritrophic interactions from historical *Bemisia* specimens. Scientific Reports 7: 429. Doi: 10.1038/s41598-017-00528-7

Tay WT, Evans GA, Boykin LM, De Barro PJ 2012. Will the Real *Bemisia tabaci* Please Stand Up? PLoS One 7: e50550. DOI: 10.1371/journal.pone.0050550

Vyskočilová S, Seal S, Colvin J 2019. Relative polyphagy of “Mediterranean” cryptic *Bemisia tabaci* whitefly species and global pest status implications. Journal of Pest Science 92: 1071–1088. doi: 10.1007/s10340-019-01113-9

Vyskočilová S, Tay WT, van Brunschot S, Seal S, Colvin J 2018. An integrative approach to discovering cryptic species within the *Bemisia tabaci* whitefly species complex. Sci Rep 8: 10886. doi: 10.1038/s41598-018-29305-w

Walsh TK, et al. 2019. Mitochondrial DNA genomes of five major *Helicoverpa* pest species from the Old and New Worlds (Lepidoptera: Noctuidae). Ecol Evol 9: 2933–2944. doi: 10.1002/ece3.4971

Wang HL, et al. 2016. First Report of Sri Lankan cassava mosaic virus Infecting Cassava in Cambodia. Plant Disease 100: 1029–1029. doi: 10.1094/Pdis-10-15-1228-Pdn

Weir BS, Cockerham CC 1984. Estimating F-Statistics for the Analysis of Population-Structure. Evol. 38: 1358–1370. doi: Doi 10.2307/2408641

Wosula EN, Chen WB, Fei ZJ, Legg JP 2017. Unravelling the Genetic Diversity among Cassava *Bemisia tabaci* Whiteflies Using NextRAD Sequencing. Genome Biology and Evolution 9: 2958–2973. doi: 10.1093/gbe/evx219

Zheng XW, et al. 2012. A high-performance computing toolset for relatedness and principal component analysis of SNP data. Bioinformatics 28: 3326–3328. doi: 10.1093/bioinformatics/bts606

